# SARS-CoV-2 binding to ACE2 triggers pericyte-mediated angiotensin-evoked cerebral capillary constriction

**DOI:** 10.1101/2021.04.01.438122

**Authors:** Chanawee Hirunpattarasilp, Gregory James, Felipe Freitas, Huma Sethi, Josef T. Kittler, Jiandong Huo, Raymond J. Owens, David Attwell

## Abstract

The SARS-CoV-2 receptor, ACE2, is found on pericytes, contractile cells enwrapping capillaries that regulate brain, heart and kidney blood flow. ACE2 converts vasoconstricting angiotensin II into vasodilating angiotensin-(1-7). In brain slices from hamster, which has an ACE2 sequence similar to human ACE2, angiotensin II alone evoked only a small capillary constriction, but evoked a large pericyte-mediated capillary constriction generated by AT1 receptors in the presence of the SARS-CoV-2 receptor binding domain (RBD). The effect of the RBD was mimicked by blocking ACE2. A mutated non-binding RBD did not potentiate constriction. A similar RBD-potentiated capillary constriction occurred in human cortical slices. This constriction reflects an RBD-induced decrease in the conversion of angiotensin II to angiotensin-(1-7). The clinically-used drug losartan inhibited the RBD-potentiated constriction. Thus AT1 receptor blockers could be protective in SARS-CoV-2 infection by reducing pericyte-mediated blood flow reductions in the brain, and perhaps the heart and kidney.

## Introduction

Despite the primary site of infection by SARS-CoV-2 being the respiratory tract, the virus evokes dysfunction of many other organs, including the brain, heart and kidney: 36% of hospitalised patients show neurological symptoms^1^, 20% develop myocardial injury^2^ and 41% experience acute kidney injury^3^. This could reflect either a spread of virus via the blood^4^, or the effects of inflammatory mediators released from the lungs. These effects may contribute to “long Covid”, in which clouding of thought and physical exhaustion extend for months after the initial infection.

The receptor^5,6^ for SARS-CoV-2 is the enzyme ACE2 (part of the renin-angiotensin system that regulates blood pressure), which converts^7^ vasoconstricting angiotensin II into vasodilating angiotensin-(1-7). The Spike protein of SARS-CoV-2 binds to ACE2 to trigger its endocytosis^6^. For the closely-related SARS virus, binding of only the receptor binding domain (RBD) is sufficient^8^ to evoke internalisation of ACE2.

In the heart^9^ and brain^10^ the main cells expressing ACE2 are pericytes enwrapping capillaries (and some endothelial cells), and pancreas and lung pericytes also express ACE2^11,12^. Pericytes express contractile proteins and in pathological conditions have been shown to decrease blood flow in the brain^13,14^, heart^15^ and kidney^16^. Interestingly, a decrease of blood flow has been reported for SARS-CoV-2 infection in the brain^17^ and kidney^18^. This could be due to pericyte dysfunction caused by SARS-CoV-2 reducing the activity of ACE2, either by occluding its binding site for angiotensin II (although this is thought not to occur for the related SARS virus^19^) or by promoting internalisation of the enzyme^6,8^. In the presence of angiotensin II (either renally-derived and reaching the brain parenchyma via a compromised blood-brain barrier, or generated by the brain’s own renin-angiotensin system^20^), a reduction of ACE2 activity would increase the concentration of vasoconstricting angiotensin II and decrease the concentration of vasodilating angiotensin-(1-7).

## Results

### Angiotensin II evokes pericyte-mediated capillary constriction via AT1 receptors

To study the effect of the SARS-CoV-2 RBD on cerebral capillary pericyte function, we employed live imaging^21^ of brain slices from Syrian golden hamsters. Hamsters have an ACE2 Spike sequence which is more similar to that in humans than is the rat and mouse ACE2 Spike sequence^22^. In particular amino acid 353 in hamsters and humans is a lysine (K) rather than a histidine (H), and this is a key determinant^23^ of how well coronaviruses bind to ACE2, making hamsters a good model for studying SARS-CoV-2 effects^22^.

We assessed the location of ACE2 and contractile properties of pericytes in the cerebral microvasculature of the hamster, which have not been studied previously (Fig. 1). Immunohistochemistry (IHC) revealed ACE2 to be predominantly expressed in capillary pericytes expressing NG2 and PDGFRβ (Fig. 1a). Quantification of overlap with the pericyte marker PDGFRβ (Fig. 1b) revealed ~75% co-localisation (Fig. 1c), and comparison of expression in capillaries and arterioles showed that capillaries exhibited ~75% of the ACE2 expression (Fig. 1d). These results are consistent with transcriptome and IHC data from mouse and human brain^10^ and human heart^9^. As for brain pericytes in rats^24^, the thromboxane A_2_ analogue U46619 (200 nM) evoked a pericyte-mediated capillary constriction and superimposed glutamate evoked a dilation (Fig. 1e).

**Fig. 1.**
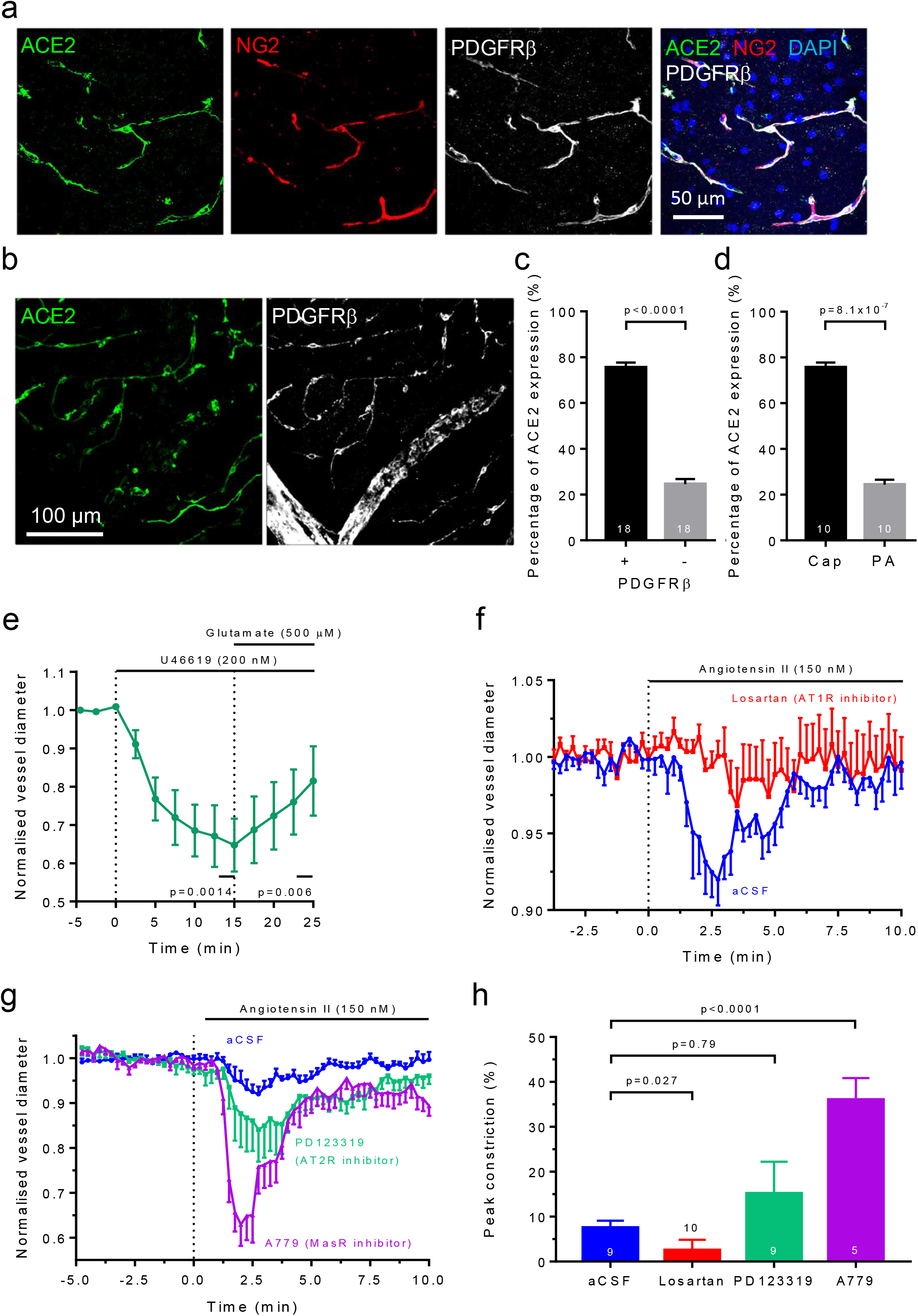
Cerebral pericytes express ACE2 and constrict capillaries in response to Ang II. (**a**) Labelling of hamster cortical slice with antibodies to the SARS-CoV-2 receptor ACE2, the pericyte markers NG2 and PDGFRβ, and with DAPI to label nuclei (**b**) Lower magnification maximum intensity projection of ACE2 and PDGFRβ labelling, showing capillaries and penetrating arteriole. (**c**) Integrated ACE2 labelling overlapping with a binarised mask of PDGFRβ labelling and with the inverse of this mask (numbers on bars are image stacks). (**d**) Integrated ACE2 labelling over capillaries versus penetrating arterioles (PA). (**e**) Average normalised diameter changes (mean±s.e.m.) at 5 pericytes in brain slices exposed to the thromboxane A_2_ analogue U46619 (200 nM), and then with the neurotransmitter glutamate (500 μM) superimposed. (**f**) Average normalised diameter changes at 9 pericytes exposed to 150 nM angiotensin II alone (aCSF), and 10 pericytes exposed to angiotensin II in the presence of the AT1R blocker losartan (20 μM). (**g**) As in (f) (aCSF plot is the same) but showing angiotensin II response in the presence of the AT_2_R blocker PD123319 (1 μM) or the Mas receptor blocker A779 (10 μM). (**h**) Peak constriction evoked by angiotensin II in different conditions (number of pericytes studied shown on bars).

Applying angiotensin II (150 nM) evoked a transient constriction, which was inhibited by the AT1 receptor blocker losartan (20 μM, Fig. 1f). Similar angiotensin-evoked pericyte-mediated capillary constriction has been reported in the kidney^25^ and retina^26^. The transience of the constriction might reflect receptor desensitisation^27^ at this relatively high angiotensin II concentration, or a delayed activation of Mas receptors after the angiotensin II is converted to angiotensin (1-7). Blocking either AT2 receptors (with 1 μM PD123319) or Mas receptors (with 10 μM A779) increased the angiotensin II evoked constriction (approximately 4.5-fold for MasR block, p<10^-4^, Fig. 1g-h), consistent with the AT1R-mediated constriction being opposed by angiotensin II activating AT2 receptors, by angiotensin-(1-7) activating Mas receptors, or by activation of AT2/Mas heteromeric^28^ receptors.

### SARS-CoV-2 binding potentiates angiotensin II evoked capillary constriction

Acute application of the RBD of Covid 19 (at 0.7 mg/l, or ~22.5 nM, which is approximately 5 times the EC_50_ for binding^29^) for up to 40 mins evoked a small and statistically insignificant reduction of capillary diameter at pericytes (Fig. 2a). On applying a very high level of angiotensin II (2 μM) in the absence of RBD, a transient constriction of capillaries at pericytes was observed (6.3±3.6% in 6 capillaries, not significantly different from the 7.5±1.6% observed using 150 nM angiotensin II in 9 capillaries in Fig. 1f, p=0.73). However, if brain slices were exposed for 30 min to RBD (0.7 mg/l) before the same concentration of angiotensin II was applied together with the RBD, then the angiotensin II evoked a 5-fold larger constriction of 31.5±9.3% in 4 capillaries (significantly different to that seen in the absence of RBD, p=0.019, Fig. 2b). The 30 min pre-exposure period was used in order to allow time for the large RBD molecule to diffuse into the slice, and was mimicked for the experiments without the RBD. This large constriction-potentiating effect of the RBD was not a non-specific effect on the pericytes’ contractile apparatus, because the contractile response to U46619 (200 nM) was unaffected by the RBD (Fig. 2c).

**Fig. 2.**
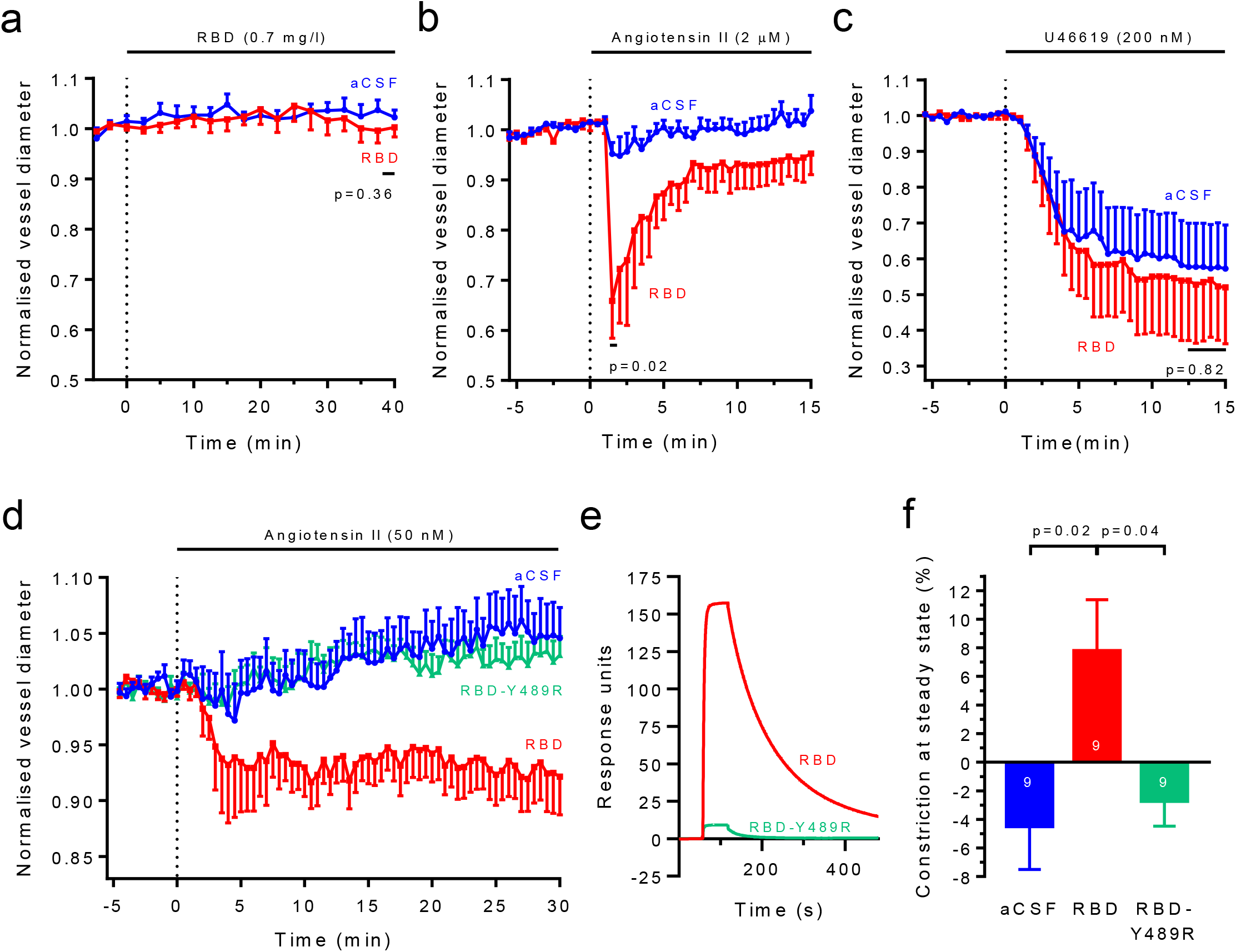
The SARS-CoV-2 RBD potentiates angiotensin-evoked capillary constriction. (**a**) Perfusion of brain slices with RBD (0.7 mg/l) has no significant effect on capillary diameter at pericytes (mean±s.e.m.; n=5 for aCSF and 6 for RBD). (**b**) After preincubation in aCSF for 30 mins, applying 2 μM angiotensin II evokes a small transient capillary constriction at pericytes (n=6), while including RBD (0.7 mg/l) in the solutions results in an ~5-fold larger response to angiotensin II (n=4; peak constriction plotted is slightly larger than the mean value quoted in the text because the latter was averaged over 5 frames and here only every 5th frame is plotted). (**c**) RBD has no effect on constriction evoked by 200 nM U46619 (n=6 for aCSF and 5 for RBD). (**d**) Response to 50 nM angiotensin II after pre-incubation and subsequent perfusion with aCSF, or aCSF containing RBD or Y489R mutant RBD (n=9 for each). (**e**) Surface plasmon resonance responses for RBD and mutant (Y489R) RBD binding to immobilised human ACE2. (**f**) Mean constriction between t = 29.67 and 30.00 min in (d).

The high concentration of angiotensin II used in Fig. 2b is probably unphysiological and evokes a transient response for reasons that are discussed above. We therefore switched to a lower angiotensin II concentration (50 nM, Fig. 2d), which is more similar to levels found physiologically within the kidney^30,31^ and heart^32^. In the presence of the RBD, the constricting response to angiotensin II was increased from an insignificant dilation of −4.5±3.0% to 7.8±3.6% (9 capillaries each, p=0.02), i.e. effectively a constriction of ~12% (from 100*{1 - (92.2%/104.5%)}).

Using surface plasmon resonance to assess binding of RBD mutants to immobilised ACE2, we identified the Y489R mutation as reducing binding by ~94% (Fig. 2e). Applying this mutated RBD (for which glycosylation of the protein is expected to be the same as for the normal RBD) had essentially no effect on the response to angiotensin II (Fig. 2d, f). Thus, the potentiation of the angiotensin II response by the RBD is a result of it binding to ACE2.

### The RBD effect is mimicked by blocking ACE2, and blocked by losartan

We hypothesised that the potentiating effect of the RBD on the response to angiotensin II reflects a decrease in the conversion by ACE2 of vasoconstricting angiotensin II into vasodilating angiotensin-(1-7). Such a decrease is expected if RBD binding promotes ACE2 internalisation^6,8^ or if it occludes the angiotensin II binding site. We therefore tested the effect of an ACE2 inhibitor (MLN-4760, 1 μM^33^) on the response to 50 nM angiotensin II. This closely mimicked the potentiating effect of the RBD, confirming that the RBD reduces effective ACE2 activity (Fig. 3a, b).

**Fig. 3.**
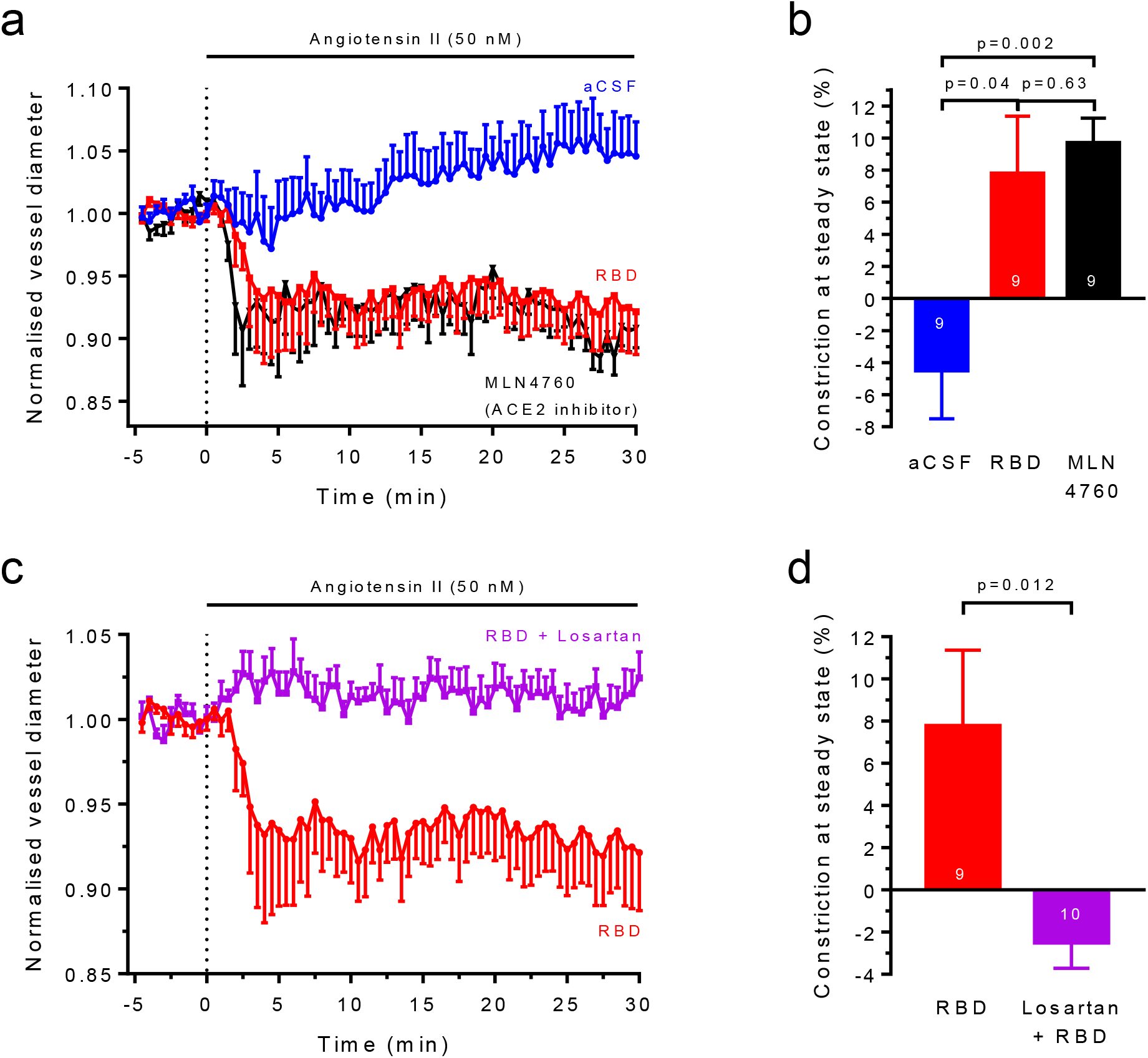
The effect of RBD is mimicked by blocking ACE2 and reduced by losartan. (**a**) Capillary constriction at pericytes in response to 50 nM angiotensin II in the absence (n=9) and presence (n=9) of the RBD (mean±s.e.m., replotted from Fig. 2d) and in the presence of the ACE2 inhibitor MLN4760 (1 μM, n=9). (**b**) Constriction in (a) between t = 29.67 and 30.00 min. (**c**) Response to 50 nM angiotensin II after 30 mins incubation in (and continued perfusion with) aCSF containing RBD (0.7 mg/l, n=9, from Fig. 2d) or additionally losartan (20 μM, n=10). (**d**) Constriction in (c) between t = 29.67 and 30.00 min.

With a view to reducing SARS-CoV-2 evoked capillary constriction and any associated reduction of microvascular blood flow, we tested whether the AT1 receptor blocker losartan prevented the constricting effect of the RBD. Losartan completely blocked the angiotensin II evoked constriction seen in the presence of the RBD (Fig. 3c, d).

In human SARS-CoV-2 infection it has been suggested that one pathological mechanism is a loss of pericytes caused by viral infection reducing their viability or their interactions with endothelial cells^10^. In a transgenic model of pericyte loss (decreasing PDGFRβ signalling) it was found that endothelial cells upregulated von Willebrand Factor (vWF) production, and thus produced a pro-thrombotic state, which could explain the coagulopathy seen in SARS-CoV-2 patients^10^. However, exposing hamster brain slices to RBD (0.7 mg/l) for 3 hours, in the absence or presence of 50 nM angiotensin II, had no significant effect on pericyte death as assessed by propidium labelling (Fig. 4a). Nevertheless, infection with the actual virus might have more profound effects on pericyte function or viability than does exposure to the RBD.

**Fig. 4.**
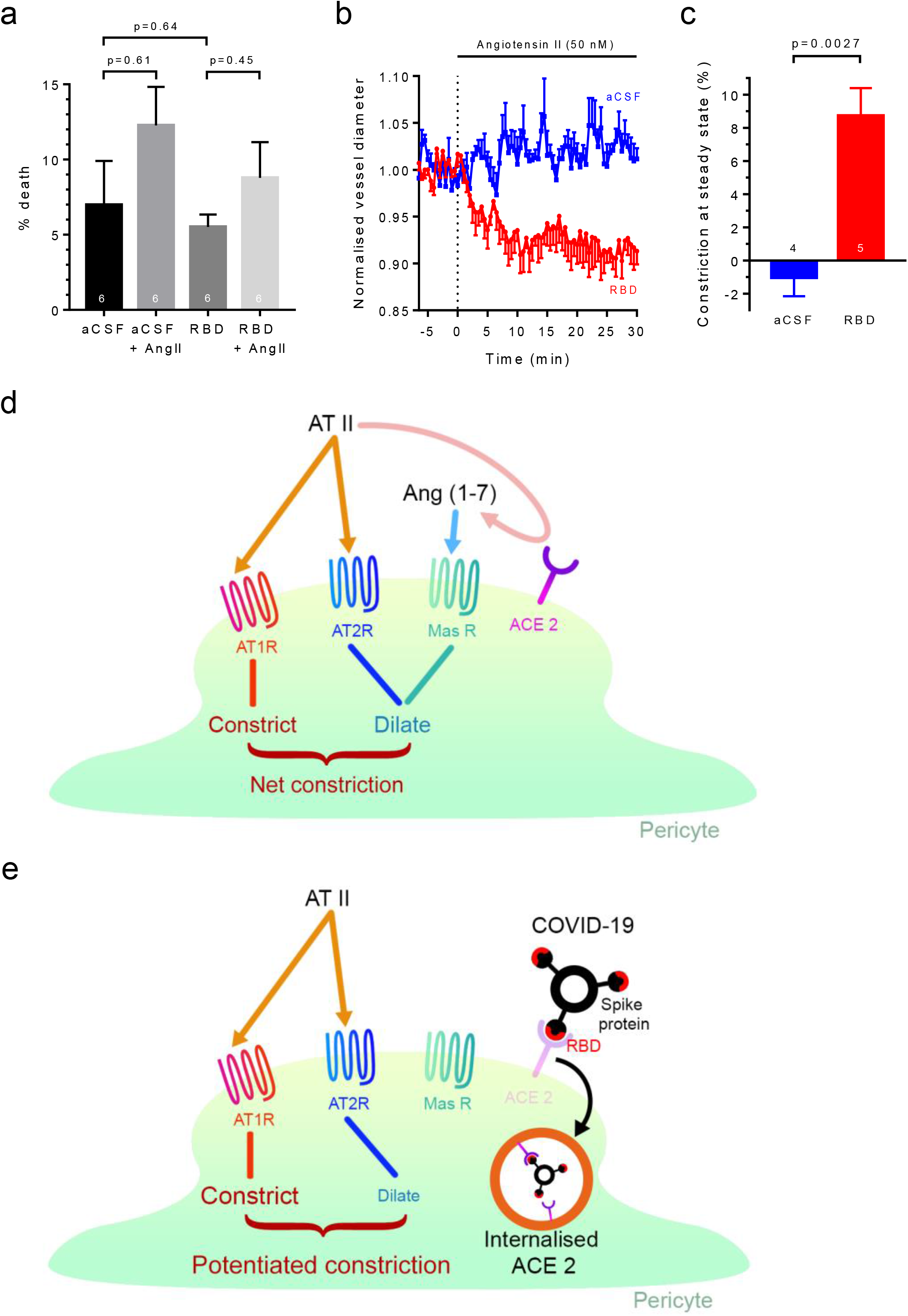
The SARS-CoV-2 RBD potentiates constriction in human capillaries. (**a**) Percentage of pericytes dead (assessed with propidium iodide) in hamster brain slices (numbers on bars) after 3 hours incubation in aCSF, or aCSF containing 50 nM angiotensin II and/or RBD (0.7 mg/l). (**b**) Effect of 50 nM angiotensin on capillary diameter (mean±s.e.m.) at pericytes in human brain slices in the presence (n=4) and absence (n=5) of the RBD (30 mins pre-incubation). (**c**) Mean constriction at 30 mins) from data in (b). (**d-e**) Likely mode of operation of RBD binding to ACE2. (**d**) Normally, angiotensin II (e.g. generated by the brain renin-angiotensin system (RAS)) can act on vasoconstricting AT_1_ receptors or vasodilating AT2 receptors, and is converted (pink arrow) by pericyte ACE2 to vasodilating angiotensin-(1-7) that acts via Mas receptors. (**e**) In the presence of SARS-CoV-2, binding of the Spike protein RBD to ACE2 leads to its internalisation (see Introduction), reducing the conversion of angiotensin II to angiotensin-(1-7). Angiotensin II (derived from the brain RAS or from the peripheral RAS) will then evoke a different balance of responses via the receptors shown, generating a larger constriction because of less activation of Mas receptors.

### Capillary constriction is potentiated by SARS-CoV-2 RBD in human capillaries

To assess whether the potentiation of capillary constriction, characterised above in hamsters, also occurs in human capillaries, we employed brain slices made from live human brain tissue that was removed in the course of tumour removal surgery^14^. Consistent with the similar binding^22^ of the SARS-CoV-2 RBD to human and hamster ACE2, we found that the RBD greatly potentiated the pericyte-mediated capillary constriction evoked by 50 nM angiotensin II (Fig. 4b-c). SARS-CoV-2 binding would therefore be expected to decrease human cerebral blood flow assuming that, as in rodents, the largest resistance to flow within the brain parenchyma is provided by capillaries^34^.

## Discussion

The data presented above are consistent with the scheme shown in Fig. 4d-e. Binding of the SARS-CoV-2 RBD to ACE2 in pericytes leads to a decrease in ACE2 activity, either as a result of ACE2 internalisation^6,8^ or due to occlusion of the angiotensin II binding site. This leads to an increase in the local concentration of vasoconstricting angiotensin II and a decrease in the concentration of vasodilating angiotensin-(1-7). The resulting activation of contraction via AT_1_ receptors in capillary pericytes reduces capillary diameter locally by ~12% when 50 nM angiotensin II is present. As most of the vascular resistance within the brain is located in capillaries^34^, this could significantly reduce cerebral blood flow (as occurs following pericyte-mediated constriction after stroke and in Alzheimer’s disease^13,14^). Presumably the same mechanism could evoke a similar reduction of blood flow in other organs where pericytes express ACE2 and AT_1_ receptors.

From our data it is complicated to predict the magnitude of blood flow reduction expected. If a 12% diameter reduction occurred uniformly in the cerebral vasculature then, by Poiseuille’s law for a pure liquid (flow proportional to the 4th power of diameter), the blood flow would be reduced by 40% [from (1-0.12)^4^ = 0.6]. However pericytes occur only every 30-100 μm (depending on age) along capillaries, suggesting a less profound effect on vascular resistance. On the other hand, the fact that blood contains cells results in its viscosity increasing dramatically at small diameters^35^, so that even small pericyte-mediated constrictions can have a large effect. Indeed, complete stalling of blood flow in capillaries can occur as a result of neutrophils (which are less distensible than red blood cells) becoming stuck at narrow parts of the vessel, for example near constricted pericytes^36–38^, and this could also transform a small constriction into a much larger reduction of blood flow.

In order for SARS-CoV-2 to evoke pericyte-mediated capillary constriction (or to cause pericyte dysfunction that upregulates vWF production^10^) the virus would need to bind to the ACE2 that is located in pericytes located on the opposite side of the endothelial cell barrier from the blood. Access to pericytes in the brain parenchyma might occur via initial infection of the nasal mucosa and movement from there up the olfactory nerve into the brain^39,40^. Alternatively, movement of the S1 part of the Spike protein across the blood-brain barrier by transcytosis has been reported^41^, and crossing the endothelial cell layer may also occur via infection of monocytes (which express ACE2 highly^42^ and can cross endothelial cells), or via breakdown of the blood-brain barrier as a result of cytokines released as a result of lung inflammation^43^.

The reduction of blood flow produced by pericyte-mediated capillary constriction, together with any upregulation of vWF that may occur^10^, will tend to promote clotting in the microvasculature. SARS-CoV-2 infection is associated with thrombus formation^44^ in large vessels that can be imaged, but it seems possible that thrombi of microvascular origin^45^ may add to this, and could perhaps even seed these larger clots. Together, capillary constriction and thrombus formation will reduce the energy supply to the brain and other organs, initiating deleterious changes that probably contribute to the long duration symptoms^46^ of “long Covid”. Indeed, the decrease of cerebral blood flow occurring during SARS-CoV-2 infection^17^ outlasts the acute symptoms^47^.

Our data suggest an obvious therapeutic approach, i.e. that the reduction of cerebral and renal blood flow that is observed in SARS-CoV-2 infection^17,18^ might be blockable using an AT_1_ receptor blocker such as losartan. Interestingly, two clinical trials (clinicaltrials.gov/ct2/show/NCT04312009 and clinicaltrials.gov/ct2/show/NCT04311177) of the possible beneficial effects of losartan in SARS-CoV-2 infection are now under way.

## Acknowledgements

Supported by a HRH Princess Chulabhorn College of Medical Science Scholarship (CH), an MRC CARP award (GJ), a Rosetrees Trust grant (CF1/100004) (DA and FF), EPSRC grant (EP/S025243/1) (RJO and JH), and ERC (BrainEnergy) and Wellcome Trust Investigator Awards (099222/Z/12/Z) (DA). RBD and mutant RBD protein were provided by the UK COVID-19 Protein Production Consortium. For the purpose of Open Access, the author has applied a CC BY public copyright licence to any Author Accepted Manuscript version arising from this submission.

## Methods

### Animals

Brain slices (200-300 μm thick) were made from the brains of Syrian golden hamsters (age 5-24 weeks) of both sexes, which were humanely killed (in accordance with UK and EU law) by cervical dislocation after being anaesthetised with isoflurane. In each slice only one pericyte was studied. The constriction evoked by angiotensin II in the presence of the RBD showed overlapping ranges of value for 2 female vessels and 9 male vessels (p=0.67).

### Imaging of pericyte mediated constriction

Pericytes on cortical capillaries were identified visually as previously described (Fig. S1 of ref. 14) and imaged with a CCD camera as described^21^. Diameter was measured in ImageJ by drawing a line across the vessel between the inner walls of the endothelial cells.

### RBD and mutant RBD synthesis

Codon optimised Genblocks (IDT Technology) for the receptor binding domain (RBD amino acids 330-532) of SARS-CoV-2 (Genbank MN908947) and human Angiotensin Converting Enzyme 2 (ACE-2, amino acids 19-615) were inserted into the vector pOPINTTGneo (PMID: 25447866) incorporating a C-terminal BirA-His6 tag and pOPINTTGneo-3C-Fc to make C-terminal fusions to Human IgG Fc. The RBD-Y489R mutant was generated by firstly amplifying the RBD-WT gene using oligos TTGneo_RBD_F and RBD-Y489R_R, as well as RBD-Y489R_F and TTGneo_RBD_R; followed by joining the two resulted fragments with TTGneo_RBD_F and TTGneo_RBD_R.

TTGneo_RBD_F 5′-gcgtagctgaaaccggcccgaatatcacaaatctttgt-3′
TTGneo_RBD_R 5′-GTGATGGTGATGTTTATTTGTACTTTTTTTCGGTCCGCACAC-3′
RBD-Y489R_F 5′-GGCGTCGAGGGTTTTAACTGTCGCTTCCCACTTCAGTCATACGG-3′
RBD-Y489R_R 5′-CCGTATGACTGAAGTGGGAAGCGACAGTTAAAACCCTCGACGCC-3′

The gene carrying the Y489R mutation was then inserted into the vector pOPINTTGneo incorporating a C-terminal His6 tag by Infusion^®^ cloning. The plasmid was sequenced to confirm that the mutation had been introduced successfully. Recombinant protein was transiently expressed in Expi293^™^ (ThermoFisher Scientific, UK) and purified from culture supernatants by an immobilised metal affinity using an automated protocol implemented on an ÄKTAxpress (GE Healthcare, UK) followed by a Superdex 200 10/300GL column, using phosphate-buffered saline (PBS) pH 7.4 buffer. Recombinant RBD-WT and ACE2-Fc were produced as described^48^. The sequence of the RBD was:

*ETG*P**N**ITNLCPFGEVF**N**ATRFASVYAWNRKRISNCVADYSVLYNSASFSTFKCYGVSPTKL NDLCFTNVYADSFVIRGDEVRQIAPGQTGKIADYNYKLPDDFTGCVIAWNSNNLDSKVGGN YNYLYRLFRKSNLKPFERDISTEIYQAGSTPCNGVEGFNCYFPLQSYGFQPTNGVGYQPYR VWLSFELLHAPATVCGPKKSTN*KHHHHHH* where the residues in italics are derived from the expression vector. Glycosylated residues are shown in bold (**N**) and the tyrosine that is mutated to arginine (Y489R) in the mutant RBD is shown underlined.

### Surface plasmon resonance

Experiments were performed using a Biacore T200 system (GE Healthcare). All assays were performed using a Sensor Chip Protein A (GE Healthcare), with a running buffer of PBS pH 7.4, supplemented with 0.005% vol/vol surfactant P20 (GE Healthcare), at 25 °C. ACE2-Fc was immobilized onto the sample flow cell of the sensor chip; the reference flow cell was left blank. RBD-WT or RBD-Y489R (0.1 μM) was injected over the two flow cells, at a flow rate of 30 μl min-1 with an association time of 60 s.

### Solutions

Brain slices were superfused at 33-36°C with solution containing (mM): 124 NaCl, 2.5 KCl, 1 MgCl_2_, 2 CaCl_2_, 1 NaH2PO_4_, 26 NaHCO_3_, 10 D-glucose and 0.1 ascorbate, bubbled with 20% O_2_/70% N2/5% CO_2_ to ensure a physiological [O_2_] was achieved in the slice^13^. The high molecular weight of the RBD (~31 kD) implies it will not diffuse rapidly from the superfusion solution into brain slices so, to apply the RBD, we pre-incubated each slice in solution containing RBD (at 35°C, to allow time for diffusion) prior to placing the slice in the imaging chamber, where it was superfused with the same solution containing the RBD. This 30 min pre-incubation time was mimicked for slices that RBD was not applied to. The same procedure was followed for the mutant RBD.

### Immunohistochemistry

Hamster brain slices were fixed in 4% paraformaldehyde (PFA) while shaking at room temperature for 20 min and washed 3 times in phosphate-buffered saline (PBS). Antigen retrieval using sodium citrate buffer (consisting of 10 mM sodium citrate, 0.05% Tween 20 and HCl to adjust the pH to 6.0) for 20 min was performed and the slices were left to cool down for 20 min before being washed in PBS for 5 minutes. Brain slices were transferred to blocking solution containing 10% horse serum, 0.2% saponin (Sigma-Aldrich, S7900), 200 mM glycine and 150 μM bovine serum albumin in PBS at 4°C, and shaken overnight. Slices were incubated in the blocking solution with primary antibodies for 72 hours at 4°C with agitation, washed with PBS 4 times, incubated with the secondary antibody overnight at 4°C with agitation and washed again 4 times with PBS. For nucleus counter staining, slices were incubated in PBS containing DAPI (100 ng/ml) for 1 h at room temperature and washed in PBS for 5 min. Primary antibodies used were goat anti-ACE2 (R&D systems, AF933, 1:200), mouse anti-NG2 (Abcam, ab50009, 1:200) and rabbit anti-PDGFRβ (Santa Cruz, sc-432, 1:200). Secondary antibodies used were Alexa fluor 488 donkey anti-goat (Invitrogen, A11055, 1:500), Alexa fluor 555 donkey anti-mouse (Invitrogen, A31570, 1:500) and Alexa fluor 647 donkey anti-rabbit (Invitrogen, A31573, 1:500).

### Pericyte death assessment

Brain slices (300 μm thick) were incubated for 3 h at 35°C in extracellular solution (bubbled with 20% O_2_, 5% CO_2_ and 75% N_2_) containing 7.5 μM propidium iodide (PI; Sigma-Aldrich, 81845) and isolectin B4 conjugated to Alexa Fluor 647 (Invitrogen, I32450) 3.3 μg/ml, with and without RBD (0.7 mg/l) and/or angiotensin II (50 nM). The slices were fixed with 4% PFA for 1 hour and washed 3 times with PBS, for 10 minutes each time. Nucleus counterstaining was achieved by incubating the slices in PBS containing DAPI (100 ng/ml) for 1 hour, washing 1 time with PBS. Imaging of Z stacks (approximately 320 μm x 320 μm x 20 μm) was performed on a confocal microscope. The first 20 μm from the surface were discarded to exclude cells killed by the slicing procedure.

### Human tissue

Live human cortical tissue was obtained from the National Hospital for Neurology and Neurosurgery (Queen Square, London). Cortical biopsies were taken from female subjects aged 40–74 undergoing tumour resection. Healthy brain tissue overlying the tumour, which would otherwise have been discarded, was used. Ethical approval was obtained (REC number 15/NW/0568, IRAS ID 180727 (v3.0), “Properties of human pericytes”, as approved on 9-10-2018 for extension and amendment) and all patients gave informed written consent. All tissue handling and storage were in accordance with the Human Tissue Act (2004).

### Statistics

Data are presented as mean±s.e.m. Data normality was assessed with Shapiro-Wilk or D’Agostino-Pearson omnibus tests. Comparisons of normally distributed data were made using 2-tailed Student’s t-tests. Equality of variance was assessed with an F test, and heteroscedastic t-tests were used if needed. Data that were not normally distributed were analysed with Mann-Whitney tests. P values were corrected for multiple comparisons using a procedure equivalent to the Holm-Bonferroni method (for N comparisons, the most significant p value is multiplied by N, the 2nd most significant by N-1, the 3rd most significant by N-2, etc.; corrected p values are significant if they are less than 0.05).

## Notes

### Competing Interest Statement

The authors have declared no competing interest.

